# Transmission Dynamics of Eastern Equine Encephalitis: Global Sensitivity Analysis and SHAP Parameter Importance in an Age-Structured Vector-Host Model

**DOI:** 10.1101/2025.01.13.632694

**Authors:** Aurod Ounsinegad, Christopher Mitchell, Nicholas Komar

## Abstract

Eastern equine encephalitis virus (EEEV) is a deadly arboviral pathogen with 30% severe case fatality. EEEV exhibits pronounced 2–3 year cyclical outbreak patterns in the northeastern United States, linked to shifts in mosquito feeding preferences between hatch-year and adult avians. We developed an age-structured vector-host model incorporating differential feeding patterns of *Culiseta melanura* mosquitoes on European Starlings and American Robins. Global sensitivity analysis revealed mosquito biting rate (*a*) as the dominant driver in transmission, with avian infectivity (*δ*) and exposure (*α*) playing secondary roles. Pairing tree-based machine learning algorithms with SHAP analysis on 100,000 parameter sets identified parameter hierarchies that govern cyclic transmission. Adult avian mortality (*µ*_*a*_) was identified as the key parameter underlying cyclic and stable transmission patterns. SHAP further revealed these patterns to be grounded on opposite sides of the same epidemiological mechanism. Stable endemics emerge from demographic stability paired with transmission optimization, while cyclic endemics emerge from demographic instability paired with minimal transmission optimization. Numerical simulations illustrated critical threshold dynamics at 0.2 ≤ *α* ≤ 0.4, where heightened hatch-year exposure triggers demographic instability responsible for observed 2–3 year cycles, while balanced exposure (*α* ≈ 0.5) leads to stable endemics. These mechanisms provide a foundation for targeted surveillance and control interventions.

## 1. Introduction

Eastern equine encephalitis virus (EEEV) is an arthropod-borne alphavirus first detected in humans in the United States in 1938, representing one of the most severe mosquito-borne pathogens in North America [3]. EEEV causes acute neuroinvasive disease with case fatality rates of approximately 30% among patients who develop severe neurological symptoms, and up to 90% in equine cases [3, 35]. The virus is maintained in an enzootic cycle primarily involving the mosquito vector, the Black-Tailed Mosquito (*Culiseta melanura*) in this paper, and avians, with occasional spillover to dead-end hosts such as humans and horses [8].

EEEV transmission exhibits distinct regional patterns across the eastern coast of the United States. In northern latitudes (e.g., Massachusetts, Connecticut, New Hampshire), surveillance data reveals pronounced 2–3 year epidemic cycles characterized by periods of intense transmission followed by apparent dormancy. Long-term surveillance in Connecticut demonstrates clear periodicity in viral activity, with mosquito abundance significantly correlated with EEEV detection (r = 0.69, p < 0.001) [2, 13]. Conversely, southern latitudes, particularly Florida, show more consistent endemic circulation with less pronounced cyclic patterns [12, 33].

Recent field studies have revealed that these transmission dynamics depend on differential mosquito feeding preferences based on the age structure of avian hosts. Throughout this study, we use young host/hatch-year avian, adult host/adult avian, and vector/mosquito interchangeably. Hatch-year avians experience 2.5–3.5 times higher infection rates than adults, creating an amplification effect during breeding seasons. Blood meal analysis reveals that *C. melanura* exhibits temporal feeding shifts, taking 26.9% of blood meals from Northern Cardinals year-round but increasing to over 40% during peak transmission periods in June [26]. This temporal synchronization between mosquito feeding behavior and avian breeding cycles provides a mechanistic explanation for observed outbreak patterns [2, 13].

The quantitative relationship between avian host demographics and EEEV transmission dynamics remains poorly understood. While previous modeling efforts have broadly examined EEEV transmission dynamics [27, 34], they have yet to explore the biological mechanisms driving the pronounced 2–3 year endemic cycles observed in northeastern states versus stable endemic patterns in southeastern states. Understanding these regional differences is critical for predicting outbreak patterns and developing targeted intervention strategies.

Mathematical modeling demonstrates that vector-host interactions can fundamentally alter transmission patterns through complex demographic processes [16, 23]. When vectors exhibit feeding preferences among host age classes, this can produce “diversity amplification” effects that concentrate transmission [23], while other configurations produce “dilution effects” that reduce transmission [16]. The complex interactions between mosquito vectors and avian hosts preceding human and equine cases exhibit various dynamics that influence regional infection patterns [10–12, 24]. Although demographic differences between young and adult avians may influence transmission [19, 29], the mechanistic drivers of regional outbreak patterns remain unclear. While quantitative analysis demonstrates that vector populations determine outbreak potential [7], the biological parameters governing whether transmission exhibits stable or cyclic dynamics have not been systematically identified.

This study develops an age-structured mathematical model combined with comprehensive sensitivity analysis and tree-based machine learning approaches to identify the biological mechanisms driving EEEV transmission patterns. We focus on the enzootic cycle involving *C. melanura* and two key avian species: European Starlings (*Sturnus vulgaris*) and American Robins (*Turdus migratorius*) [18]. Using Sobol global sensitivity analysis and SHAP feature importance applied to machine learning analysis of 100,000 biologically plausible parameter combinations, we identify distinct biological mechanisms governing transmission potential versus dynamical stability in EEEV. Our findings reveal that transmission efficiency parameters, particularly mosquito biting rate and the avian exposure coefficient, control outbreak potential, while demographic parameters such as adult avian mortality serve as primary drivers of cyclic versus stable dynamics. Model simulations validate these analytical findings by reproducing the three transmission regimes observed across the eastern United States. The resulting age-structured framework provides mechanistic insight into EEEV outbreak patterns and identifies key biological targets for intervention.

## 2. Model Development

In developing the stage-structured model, the host avian class is split into hatch-year avians (*y*) and adult avians (*a*) to better understand host age-specific transmission dynamics where both avian classes contain three stages: susceptible (*S*), infected (*I*), and recovered (*R*) (Fig. 1). Both classes have recovered stages where they are classified as non-competent hosts, *R*_*y*_ (Eq. 2.3) and *R*_*a*_ (Eq. 2.6), respectively, implying an inability for transmission. For this model, it is assumed that once an individual from either avian class enters the recovered stage, they remain recovered for life. This is assumed as both avian species considered for this study maintain recovery status for the same amount of time as their average lifespan [17, 25].

**Fig. 1:**
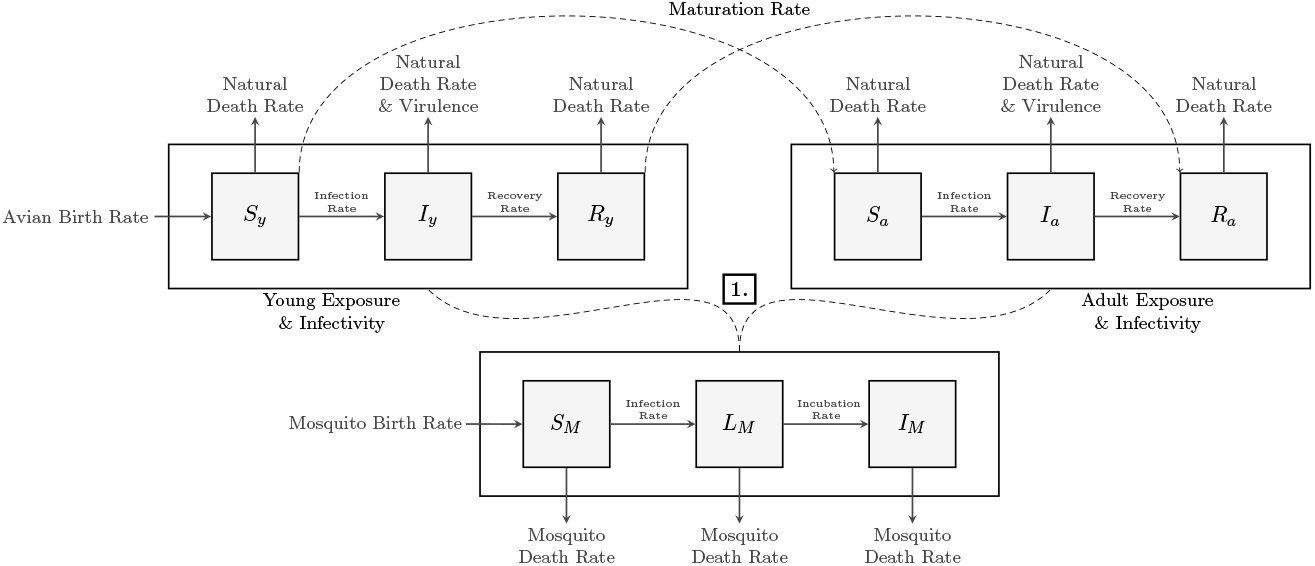
EEEV stage-structured model flow chart. The top left, top right, and bottom center boxes represent hatch-year (*y*), adult avian (*a*), and mosquito (*M*) classes, respectively. Solid directional arrows connect stages within each class, indicate their interactions, and show entrance and exit into the system. Dashed directional arrows show maturation between young and adult avian classes, where it is assumed young hosts recover before maturing. Dashed lines between vectors and hosts show bidirectional viral transmission, where **1**. represents feeding success (successful viral transmission) and host preference.

Young hosts mature into adults at a rate *m*_*y*_ from the susceptible and recovered stages, but not the infected stage. It is assumed that young hosts must recover and enter the recovered stage before they can mature. It is also assumed that there is a constant birth rate, *ψ*_*B*_, within the avian population paired with constant natural mortality rates for both hosts in each stage given by *µ*_*y*_ and *µ*_*a*_, respectively. It should be noted that the carrying capacity term, 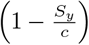, keeps logistic growth of the avian population within the model while simultaneously keeping the population bounded, where *c* denotes the numerical carrying capacity.

Through the development of the vector transmission dynamics within this model, it was determined that the mosquito class has a different stage structure compared to that of the avian host; the mosquito class moves between a susceptible (*S*), latent (*L*), and infected (*I*) stage. It is assumed that all mosquitoes are recruited into the susceptible class, *S*_*M*_ (Eq. 2.7), and that a vector must move from the latent stage, *L*_*M*_ (Eq. 2.8), into the infected stage, *I*_*M*_ (Eq. 2.9), before they are considered infectious. This is because an extrinsic incubation period must pass before infection can take place [30, 34]. Additionally, once a mosquito has made it to the infectious stage, it is assumed that it remains infectious for life. A constant birth rate, *ψ*_*M*_, and mortality rate, *µ*_*M*_ in and out of the mosquito class is also assumed.

It was observed that the relative number of mosquito bites on each avian species is determined by the number of female mosquitoes present and the abundance of avian hosts within the model [24]. However, the value for the biting rate parameter, *a* is simplified with this model, whereas the realistic transmission of EEEV occurs between a variety of different host and vector species.

Mosquito bites are distributed among both the young and adult host classes, where the exposure coefficient of the hosts to mosquito bites is given by 1 − *α* and *α*, respectively. If *α* ≈ 0.5, then each avian class is bitten in proportion to the abundance of its respective population, and all avians receive a similar number of bites. Higher *α* values correlate with increased adult host exposure, whereas lower *α* values correlate with increased young host exposure.

Susceptible hosts, *S*_*y*_ (Eq. 2.1) and *S*_*a*_ (Eq. 2.4), can move into their respective infectious class, *I*_*y*_ (Eq. 2.2) and *I*_*a*_ (Eq. 2.5), once bitten by an infectious mosquito vector, *I*_*M*_, dependent on their exposure to the virus (*α*). Although it has been observed that EEEV can be transmitted vertically within the avian population [30], vertical transmission is ignored within the model, and the principal method of transmission is observed to occur between vectors and hosts during the viral amplification season [19, 22]. Transmission of the virus from infectious mosquito vectors to susceptible hosts is partially determined by the proportion of susceptible hosts to the total population of avians 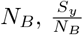 and 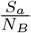 . The class-dependent virulence is given by *ν*_*y*_ and *ν*_*a*_ and the class-dependent recovery rate, is given by *γ*_*y*_ and *γ*_*a*_. To analyze the independent effect of differential avian class viral exposure, it is assumed that infectivity and recovery rate baseline values of each avian class are equal.

Mosquitoes within the susceptible stage can move into the latent stage, *L*_*M*_, after successfully biting an infectious host. Transmission of the virus from infectious avian hosts to susceptible mosquito vectors is partially determined by the proportion of susceptible mosquitoes to the total population of mosquitoes 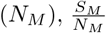. Susceptible mosquitoes take *a* bites per day, with respect to both host classes, where only a fraction of the bites taken are from infectious hosts, 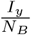 and 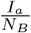. The probability that vector-host transmission will successfully occur per bite, also known as avian infectivity, is given by *δ*. Latent mosquitoes move into the infectious stage at a rate *k*, where 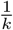 is the extrinsic incubation period of the virus that must pass before a mosquito can be considered fully infectious and transmit the virus. Although it has been observed that EEEV can affect the reproductive fitness of *C. melanura* [6], due to short life span, it is assumed that infectious mosquitoes do not have a virulence, never recover, and can only exit the system through natural death.

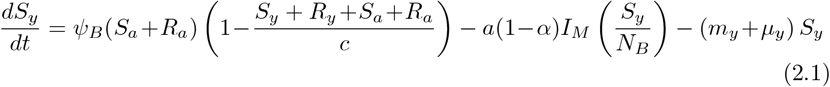

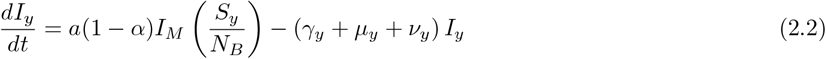

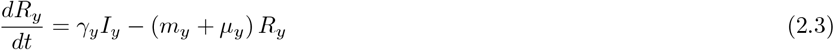

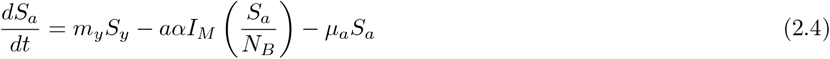

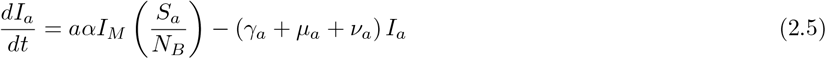

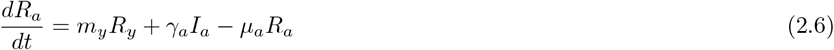

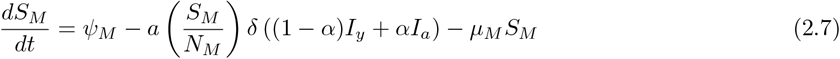

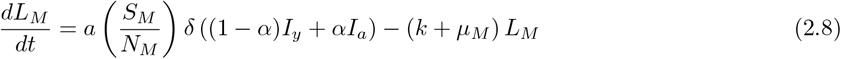

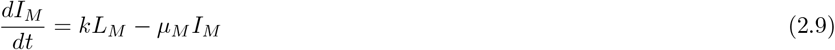

All model parameters are assumed positive. The system of ordinary differential equations the model is comprised of is presented above, with complete parameter descriptions and values in Table 1.

**Table 1:** EEEV stage-structured model parameter list. Note that total population counts for avians and mosquitoes (*N*_*B*_ and *N*_*M*_, respectively) don’t have references due to their constant change in model simulations.

| Parameter | Description | Value | Reference |
| --- | --- | --- | --- |
| $\psi_B$ | Hatch-year avian birth rate | 0.05 avians day<sup>-1</sup>/avians | [17, 25] |
| $a$ | Mosquito biting rate | 2.5 day<sup>-1</sup> | [24] |
| $\alpha$ | Avian exposure coefficient | $0 \leq \alpha \leq 0.5$ | Varied |
| $\gamma_y$ | Hatch-year avian recovery rate | $\frac{1}{3}$ day<sup>-1</sup> | [19] |
| $\gamma_a$ | Adult avian recovery rate | $\frac{1}{3}$ day<sup>-1</sup> | [19] |
| $\mu_y$ | Hatch-year avian death rate | 0.002 day<sup>-1</sup> | [17, 25] |
| $\mu_a$ | Adult avian death rate | $\frac{1}{547}$ day<sup>-1</sup> | [17, 25] |
| $\mu_M$ | Mosquito death rate | 0.1 day<sup>-1</sup> | [6] |
| $\nu_y$ | Hatch-year avian virulence | 0.5 day<sup>-1</sup> | [22] |
| $\nu_a$ | Adult avian virulence | 0.5 day<sup>-1</sup> | [22] |
| $\psi_M$ | Mosquito birth rate | 3,000 mosquitoes day<sup>-1</sup> | [6] |
| $\delta$ | Avian infectivity | 0.2 | [19] |
| $k$ | Extrinsic viral incubation period | $\frac{1}{3}$ day<sup>-1</sup> | [30] |
| $m_y$ | Avian maturation rate | 0.04 day<sup>-1</sup> | [32] |
| $N_B$ | Total population of avians | Varies | |
| $N_M$ | Total population of mosquitoes | Varies | |
| $c$ | Avian population carrying capacity | Varies | [17, 25] |

Susceptible young hosts (*S*_*y*_) are recruited at a rate of *ψ*_*B*_, where it is assumed that a majority of female avians are capable of laying two clutches of eggs per year with a mean clutch size of five [17, 25]. To bound the growth of the avian population, a carrying capacity *c* is used, where it is assumed that the carrying capacity of the avian population will remain constant.

## 3. Sensitivity Analysis

A comprehensive sensitivity analysis was conducted to identify and quantify the relative influence of model parameters on peak infection levels across three key populations: hatch-year avians (*I*_*y*_), adult avians (*I*_*a*_), and mosquitoes (*I*_*M*_). The basic reproduction number ℛ_0_ was derived using the next generation matrix approach:

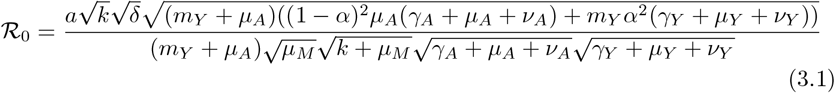

### 3.1. Parameter Sensitivity Patterns and Mechanistic Interpretation

Sobol sensitivity analysis is a variance-based global sensitivity method that quantifies the contribution of each input parameter to the total variance in model outputs [31]. The method decomposes output variance into components attributable to individual parameters (first-order indices, S1) and their interactions with other parameters (total-effect indices, ST). First-order indices measure the direct effect of a parameter when varied independently, while total-effect indices capture both direct effects and all interactions involving that parameter. This approach is particularly valuable for complex biological models where parameter interactions may be substantial, as it provides model-independent measures of parameter importance across the entire parameter space rather than at a single point.

Both local (Fig. 2) and global sensitivity (Fig. 3) analyses demonstrate that mosquito biting rate (*a*) emerges as the most important parameter across all model outputs, with Sobol total-effect indices (ST) ranging from 0.368 to 0.491. This dominance validates the central role of vector-host contact frequency in arboviral transmission [4, 24]. Avian infectivity (*δ*) consistently ranks second (ST values 0.120–0.231), matching patterns reported for other vector-borne diseases where transmission efficiency parameters dominate system behavior [29].

**Fig. 2:**
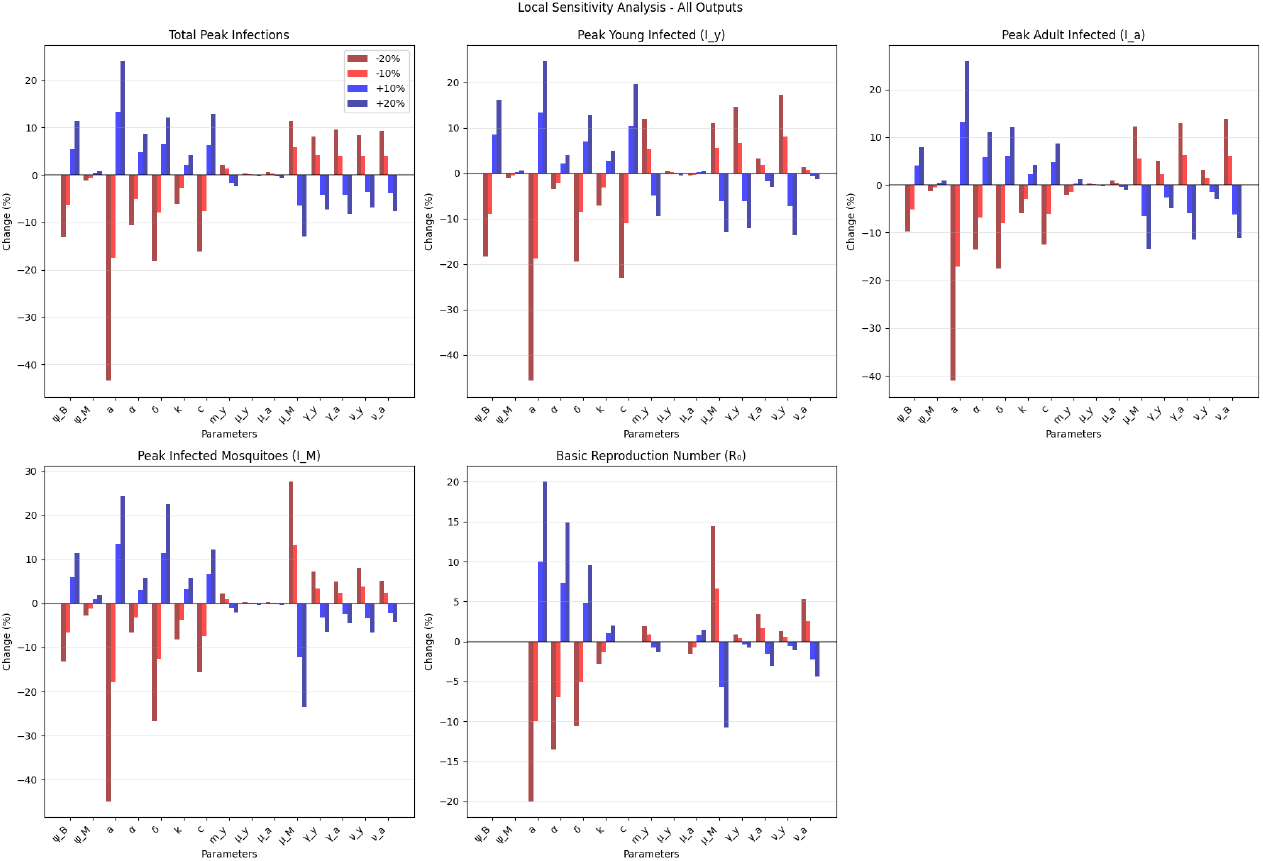
Local sensitivity analysis for EEEV transmission showing percentage changes in model outputs when parameters are varied by ±10% and ±20%. **Top row:** Total peak infections, peak young infected (*I*_*y*_), and peak adult infected (*I*_*a*_). **Bottom row:** Peak infected mosquitoes (*I*_*M*_) and basic reproduction number (ℛ_0_). Red bars indicate negative parameter effects while blue bars indicate positive effects.

**Fig. 3:**
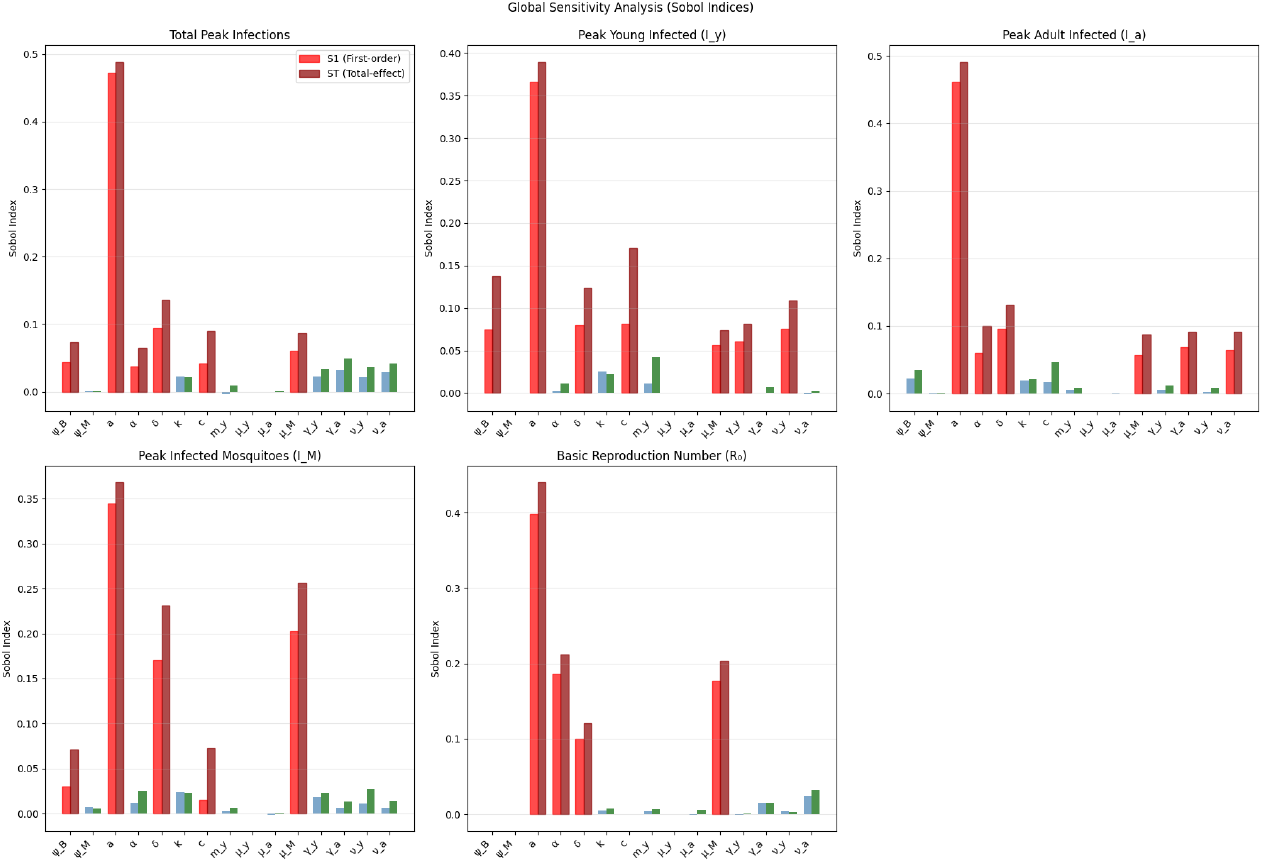
Global sensitivity analysis using Sobol indices for EEEV transmission. Red bars represent the five highest paramters indices, while blue/green are lower values. Shows first-order effects (S1: light-red) and total effects (ST: dark-red) for each parameter across multiple model outputs. Higher Sobol index values indicate greater parameter importance in determining model output variability.

The parameter hierarchy shows agreement between local and global sensitivity analyses for the top parameters (*a, δ, µ*_*M*_), confirming that parameter influences reflect genuine biological processes rather than methodological artifacts. The influence of demographic parameters (*ST* < 0.11) compared to transmission parameters validates that EEEV outbreaks are driven by transmission efficiency rather than virulence alone, consistent with field observations [10].

### 3.2. Age-Structured Population Responses and Vector Dynamics

Different sensitivity patterns between hatch-year and adult avian populations validate our age-structured approach. Both avian classes show strong sensitivity to mosquito biting rate, with adults being more responsive (ST = 0.491) than young avians (ST = 0.390). However, hatch-year avians show unique sensitivity to carrying capacity (*c*; ST = 0.171) while adults show minimal response (ST = 0.046).

The avian exposure coefficient (*α*) reveals important age-specific patterns: young avians show minimal sensitivity (ST = 0.011), adults demonstrate moderate sensitivity (ST = 0.100), and *α* strongly influences ℛ_0_ (ST = 0.212). This pattern indicates that differential exposure between age classes becomes critical for determining overall transmission potential, validating biological evidence that mosquito feeding preferences significantly alter outbreak dynamics [24].

For vector populations, while mosquito biting rate maintains dominance (ST = 0.368), the mosquito death rate emerges as the second most influential parameter (ST = 0.257), validating the critical importance of vector longevity for EEEV transmission. This confirms that vectors must survive long enough for viral development and subsequent transmission, aligning with field observations that environmental factors affecting mosquito survival strongly influence outbreak probability [7, 26].

## 4. Machine Learning Analysis and Parameter Importance

To quantify parameter interactions and identify biological mechanisms driving different endemic equilibria, we employed tree-based machine learning algorithms on 100,000 biologically plausible parameter sets. Based on epidemiological observations of transmission patterns across different regions [2,7], we classified endemic equilibria into two categories: Stable Endemic Equilibria (Stable EE), seen as persistent non-oscillatory transmission comprising ∼ 98.5% of all endemic equilibria; and Unstable Endemic Equilibria (Unstable EE), seen as rare cyclic dynamics with 2–3 year periodicity comprising ∼ 1.5% of all endemic equilibria. The extreme class imbalance between stable and unstable EE reflects the rarity of cyclic EEEV behavior.

Parameter correlation analysis showed very low correlation and confirmed independence between all biological parameters (0.0 ≤ |*r*| ≤ 0.25), validating that each parameter contributes unique information without confounding analyses (Fig. 4).

**Fig. 4:**
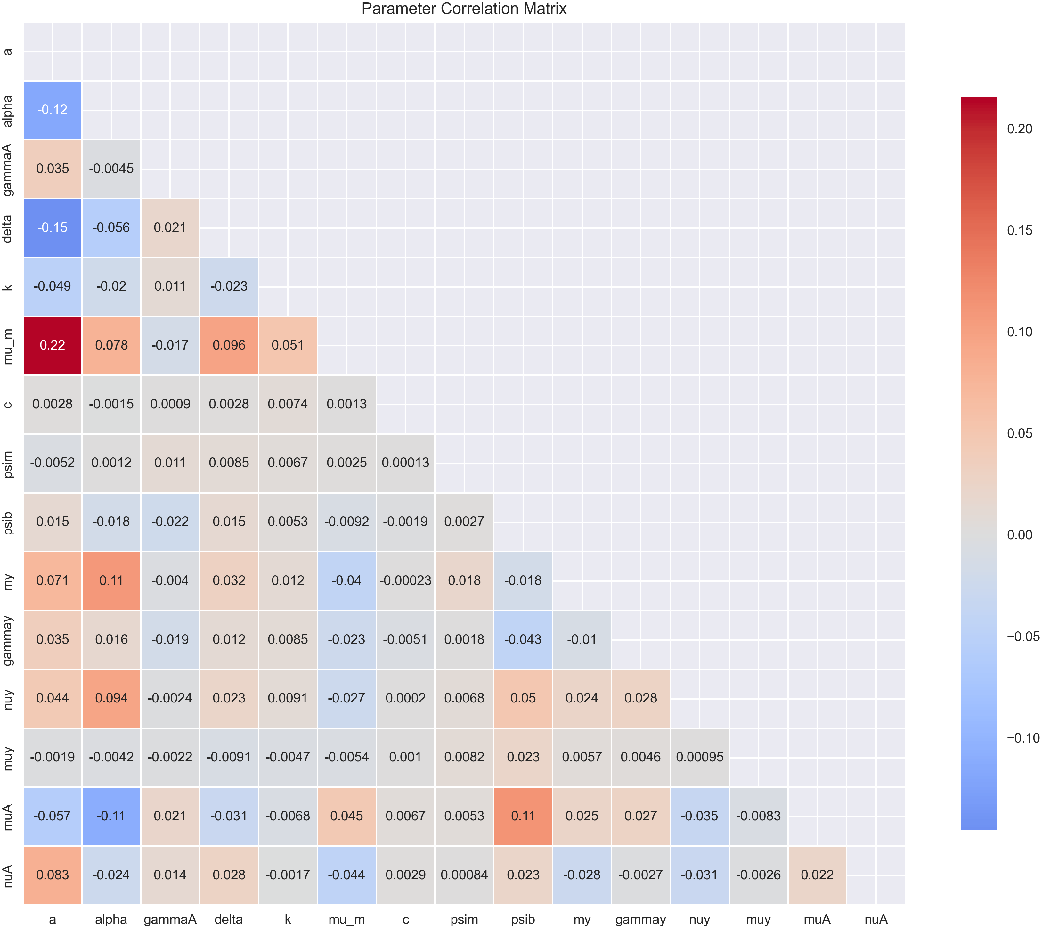
Parameter correlation matrix showing very low correlation values across all parameters (0.0 ≤ |*r*| ≤ 0.25), indicating negligible collinearity between parameters, supporting independent parameter influence on system behavior. Grey to blue boxes show negative correlations, and orange to red boxes show positive correlations.

### 4.1. Methodology and Model Performance

Three machine learning algorithms were chosen for their ability to capture non-linear parameter interactions while providing interpretable parameter importance measures: Random Forests (RF) [5, 14], eXtreme Gradient Boosted trees (XGB) [9], and Light Gradient Boosted Machine (LGBM) [15]. Tree-based algorithms excel at modeling threshold effects and parameter interactions characterizing biological systems, making them ideal for understanding EEEV’s complex transmission dynamics [21].

All models were optimized using Optuna Bayesian hyperparameter tuning and 5-fold stratified cross-validation over 30 trials to maintain proper class distribution [1, 28]. Particularly, due to the extreme class imbalance, we utilized Area Under the Precision-Recall Curve (AUPRC) in addition to Area Under the Receiver Operating Curve (AUROC) metrics to track any indication of overfitting. AUROC metrics can be misleading when there is an extreme class imbalance present in the data, which is where AUPRC becomes a more desirable performance metric.

With unstable EE set as the target variable, all models overfit due to class imbalance, with LGBM overfitting least, showing better overall generalization (Fig. 5).

**Fig. 5:**
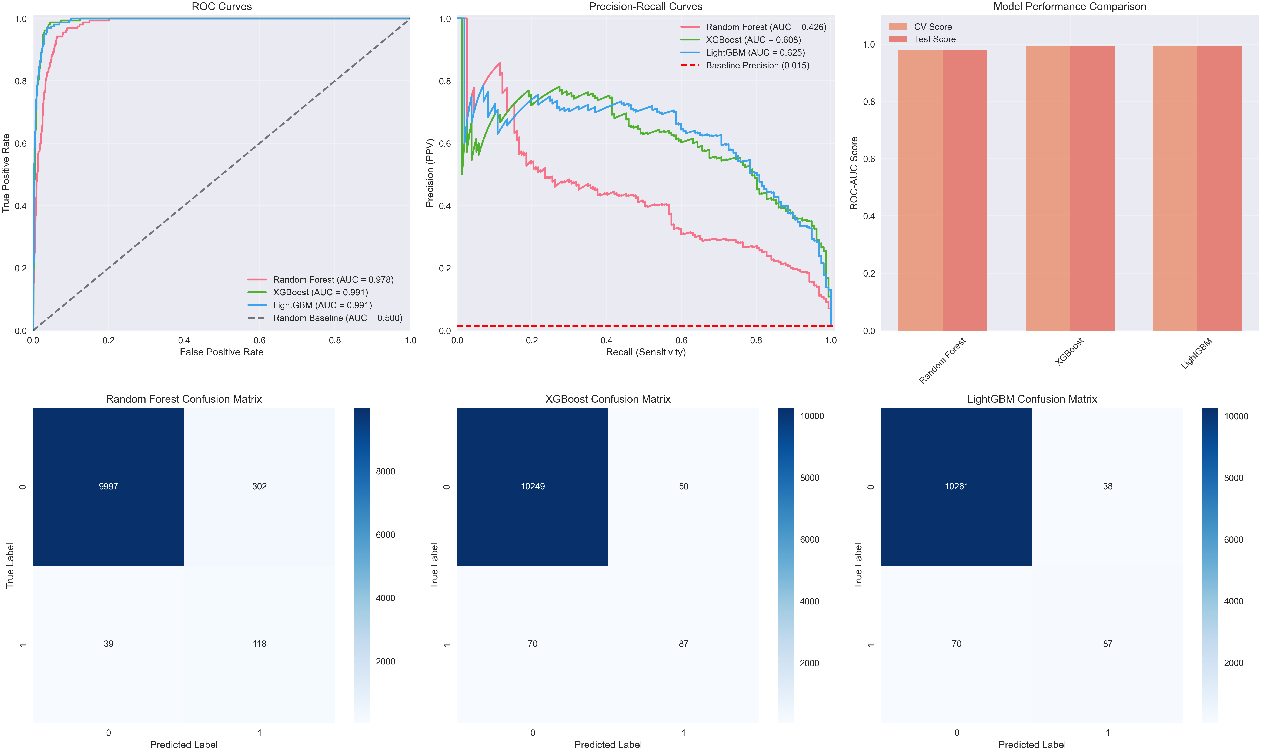
Model performance comparison: AUROC (top right), AUPRC (top center), AUROC Bar Plot comparison between cross-validated data and test data performance (top left), and Confusion Matrices (bottom row of plots; from right to left: RF, XGB, and LGBM). Given that all models overfit due to class imbalance, LGBM performed the best (AUROC: 0.991; AUPRC: 0.625), followed by XGB (AUROC: 0.991; AUPRC: 0.608), and lastly RF (AUROC: 0.978; AUPRC: 0.426).

### 4.2. Shapley Value Parameter Importance Analysis

SHAP (SHapley Additive exPlanations) value analysis, a model-agnostic parameter importance algorithm based on game theory [20], revealed distinct mechanistic drivers for both types of endemic equilibria. SHAP values offer local explanations for individual predictions and global parameter importance rankings, enabling comprehensive understandings of how each parameter drives endemic behavior across the parameter space. To allow for meaningful comparison across models, SHAP was normalized by dividing each feature’s SHAP value by the maximum absolute value within that model. This normalization approach preserves the relative importance rankings while standardizing the scale across models: 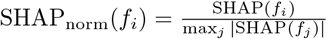, where *f*_*i*_ represents the *i*-th feature. This ensures parameter importance rankings reflect biological influence rather than model-specific architectures.

One common way to visualize SHAP values is through the use of SHAP Summary Plots (Fig. 6). In summary plots, each point represents one case (e.g., an unstable or stable EE), and each case will have exactly one point for each of the parameters shown on the y-axis. As can be seen in Fig. 6, the color of the points represents the value for each respective parameter, with blue (cooler colors) being low parameter values, purple (warmer colors) being mid parameter values, and pink (hottest colors) being high parameter values. Lastly, the x-axis shows the SHAP value for each respective point, with negative SHAP values representing a negative impact on the model’s output (e.g., the parameter is pushing the model to predict “Stable EE” for the respective case) and positive SHAP values representing a positive impact on the model’s output (e.g., the parameter is pushing the model to predict “Unstable EE” for the respective case). The combination of the color of each point (parameter value) and the respective SHAP value (negative/positive) in the plot provides information about whether a parameter has negative/positive/no correlation with model output.

**Fig. 6:**
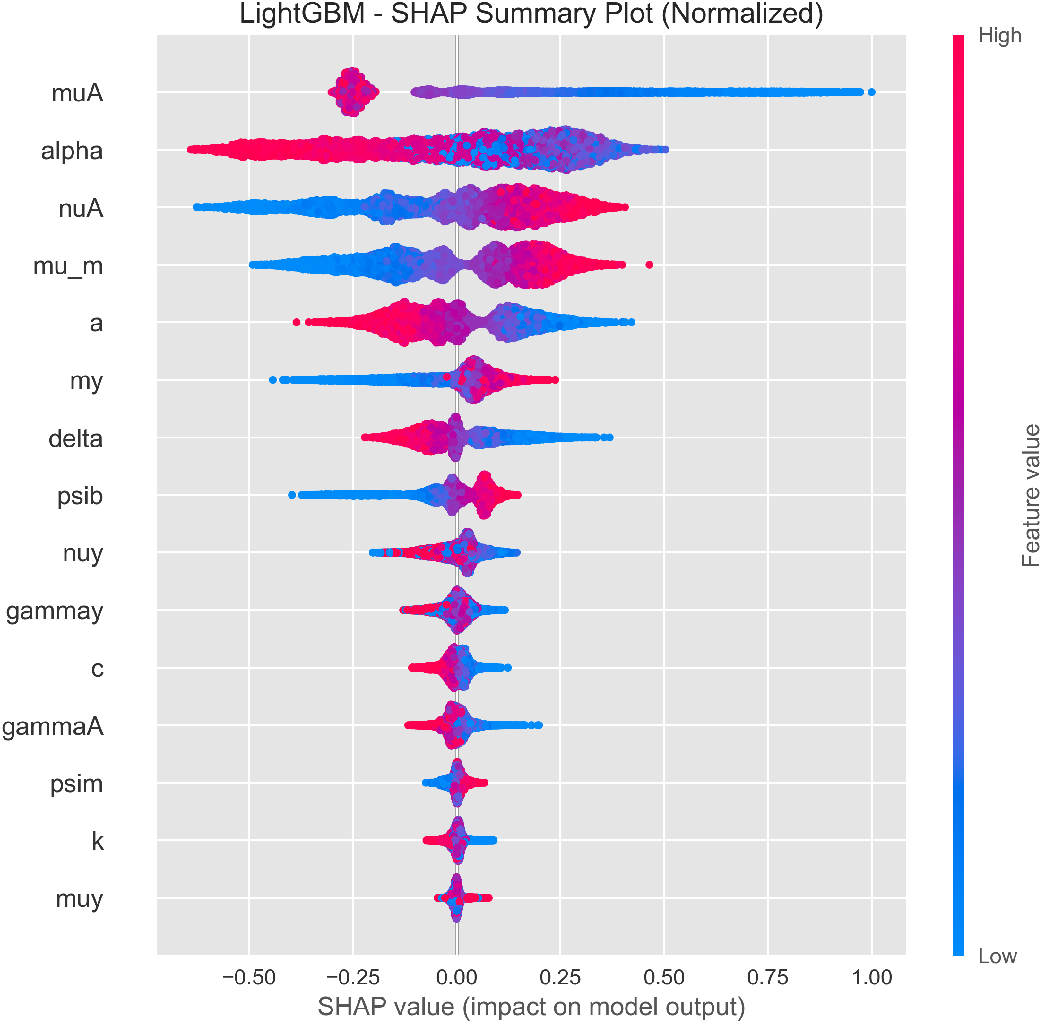
Unstable EE LGBM Model SHAP Summary Plot showing SHAP value distributions for all parameters with *µ*_*a*_, *α*, and *ν*_*a*_ ranking highest (parameter importance is read from top to bottom). Color gradients indicate parameter values with blue (cooler colors) being low, purple (warmer colors) being mid, and pink (hottest colors) being high parameter values. Positive/Negative SHAP values represent positive/negative impact on the model’s output, respectively. The combination of each point’s color and SHAP value provides information about whether a parameter has negative/positive/no correlation with model output. **Note:** The color gradient label, “Feature Value”, is the same as “Parameter Value”.

Demographic instability mechanisms tended to dominate across all models with the adult death rate (*µ*_*a*_), avian exposure coefficient (*α*), adult virulence (*ν*_*a*_), mosquito death rate (*µ*_*m*_), and mosquito biting rate (*a*) surfacing as the top five most important parameters in order (Fig. 6 shows this in the best-performing model, LGBM). *µ*_*a*_, *α*, and *a* all show a negative correlation between parameter and SHAP values with average absolute SHAP values of ∼ 0.455, 0.221, and 0.107, respectively. *ν*_*a*_ and *µ*_*m*_; however, show clear positive correlation between parameter and SHAP values with average absolute SHAP values of ∼ 0.182 and 0.140 respectively.

This hierarchy uncovers two main drivers in both types of endemic equilibria. First, with *µ*_*a*_ achieving importance values up to twice those of the second most important parameter, this shows that adult avian population stability in the system is crucial in maintaining cyclic transmission patterns. Second, *α* exhibits a more complex negative correlation than *µ*_*a*_ or *a*, demonstrating that higher exposure in adults (*α* ≥ 0.7) results in more stable EE; however, when exposure begins to favor young avians around the mid-to-low range (0.2 ≤ *α* ≤ 0.4), you see more unstable EE. Overall, the hierarchy of parameter importance indicates that cyclic dynamics emerge more through population-level demographics rather than transmission optimization.

The SHAP ridge plot further reveals the underlying distribution of the demographic instability mechanism through a fine-tuned balance between population-level parameters—parameters that can be modulated through habitat conservation and demographic support during vulnerable periods (Fig. 7 shows this in the best-performing model, LGBM).

**Fig. 7:**
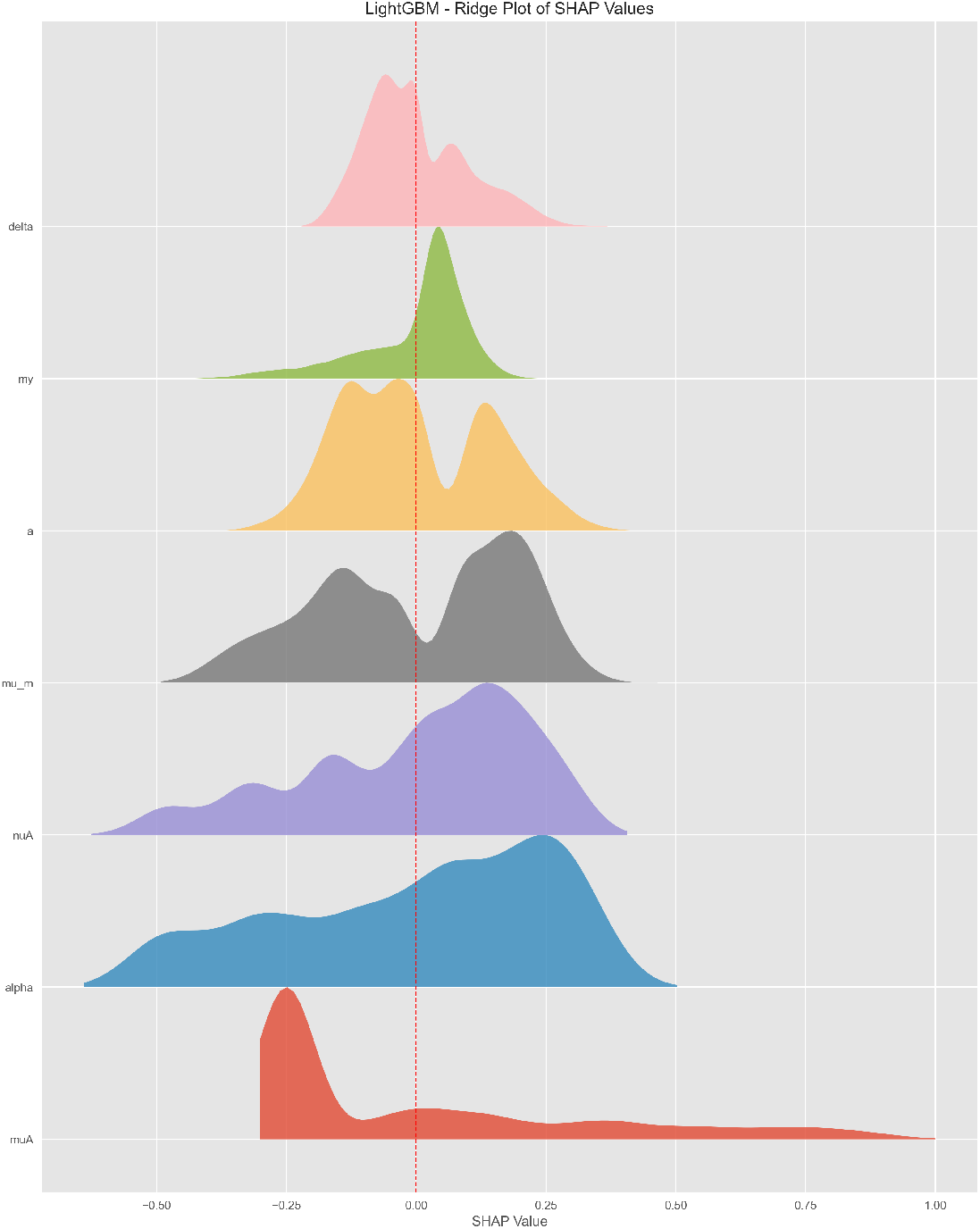
Unstable EE LGBM Model SHAP Ridge Plot comparison showing SHAP value distributions for all model parameters with top parameters *µ*_*a*_, *α*, and *ν*_*a*_ (parameter importance is read from bottom to top—opposite of Fig. 6). Each parameter’s respective ridge plot shows the broad distribution of SHAP values, with higher ridges indicating a greater frequency of the respective SHAP value.

### 4.3. Biological Validation and Intervention Implications

The unique parameter importance hierarchy shows that stable and unstable EE are both grounded in the same epidemiological demographic mechanism, but in fundamentally different ways. Stable EE depends more on demographic stability and transmission optimization through higher mosquito biting rates as well as higher natural death and exposure rates in adult avians paired with lower death in mosquitoes and virulence in adult avians (reflecting true transmission patterns in the southeastern United States) [33]. Unstable EE depends on the polar opposite: demographic instability and minimal transmission optimization through lower natural death, but higher virulence, in adults, higher exposure in young avians, and higher death, but lower biting rates, in mosquitoes (reflecting the true complex transmission patterns in the northeastern United States) [2, 13].

Mann-Whitney U-tests revealed significant parameter distribution differences (*p* < 0.001) between stable and unstable EE for over half the model parameters. Sensitivity and SHAP analysis approaches reveal complementary yet distinct parameter importance hierarchies. Sobol sensitivity analysis consistently identifies mosquito biting rate (*a*) as the most dominant parameter across model outputs (*ST >* 0.4), reflecting its control over transmission intensity. In contrast, SHAP analysis highlights demographic parameters as primary drivers of model behavior, with adult mortality (*µ*_*a*_) showing the highest parameter importance (SHAP = 0.455), followed by exposure coefficient (*α*; SHAP = 0.221) and adult virulence (*ν*_*a*_; SHAP = 0.182). Both methods generally agreed on the critical importance of the exposure coefficient, though they emphasized different aspects of its influence on system dynamics. This methodological divergence suggests that global sensitivity analysis captures transmission threshold effects while machine learning approaches emphasize parameters governing dynamical stability and oscillatory behavior within endemic states.

Since both types of endemic equilibria are grounded in the same epidemiological demographic mechanism, targeted intervention strategies are quite straightforward: emphasize host population stability (habitat conservation, disease surveillance, focus on adult avian health, etc.). The rarity of unstable EE (1.5% of endemic equilibria) reflects the biological reality that cyclic transmission represents exceptional rather than typical EEEV behavior.

## 5. Model Simulations and Results

The host stage-structured EEEV model was implemented using parameter values from Table 1 with systematic variation of key parameters identified through sensitivity and data analysis. Model simulations validate both the theoretical ℛ_0_ framework and parameter importance hierarchies revealed through global sensitivity analysis and machine learning approaches.

Numerical simulations confirm that the calculated ℛ_0_ (Eq. 3.1) accurately predicts transmission outcomes across the parameter space. Fig. 8 demonstrates how the avian exposure coefficient (*α*) interacts with key transmission parameters identified in previous sections. Panels (A) and (C) show nearly vertical ℛ_0_ = 1 boundaries, indicating that the exposure coefficient dominates transmission viability over both adult avian virulence (*ν*_*a*_) and mortality (*µ*_*a*_). Panel (B) reveals that higher mosquito mortality (*µ*_*M*_) requires a correspondingly higher exposure coefficient to maintain ℛ_0_ *>* 1, confirming vector survival as a critical constraint. Panel (D) shows that despite wide variation in mosquito biting rate (*a*), viable transmission requires exposure coefficients within a narrow window, creating a steep threshold that validates the importance of balanced age-structured exposure. Panel (E) demonstrates that avian infectivity (*δ*) operates within an exceedingly narrow range, with little influence from the exposure coefficient, suggesting a limited threshold effect despite its secondary importance in sensitivity analyses.

**Fig. 8:**
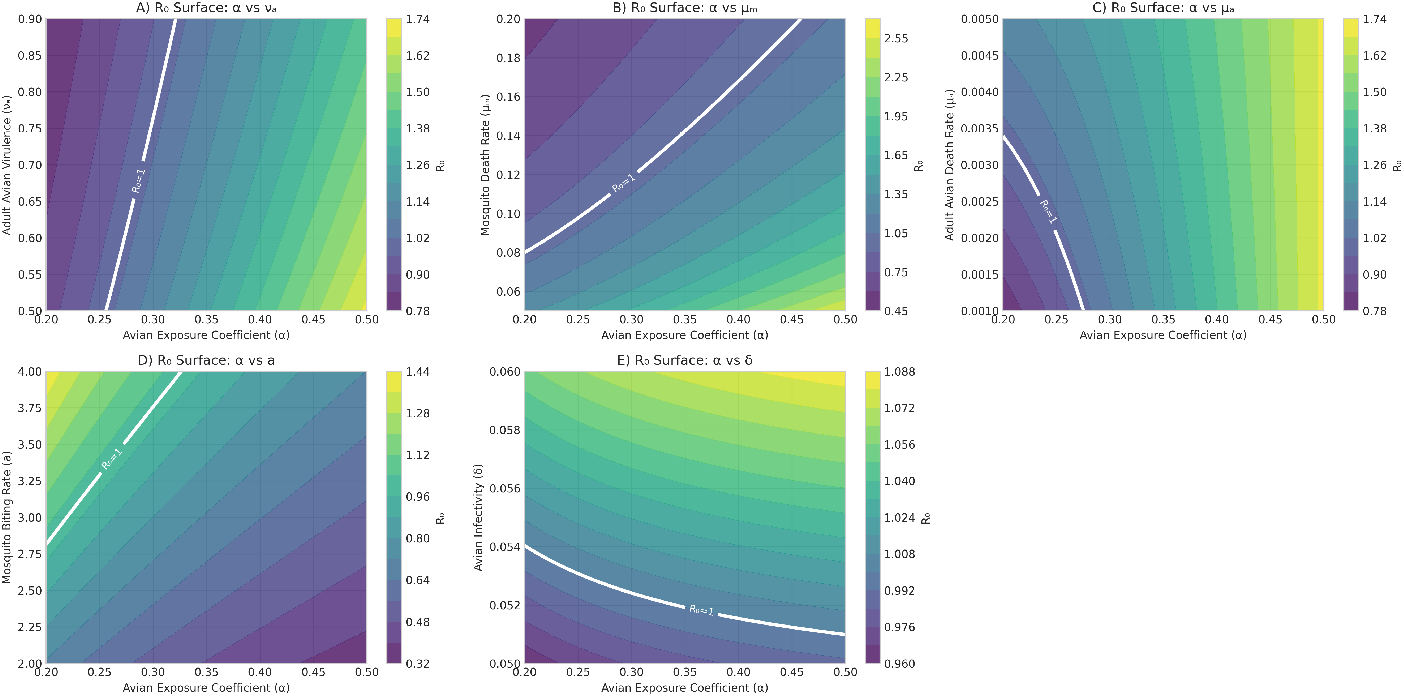
ℛ_0_ heatmap showing interactions between avian exposure coefficient (*α*) and key transmission parameters. **(A)** Adult avian virulence (*ν*_*a*_) versus exposure coefficient showing demographic effects. **(B)** Mosquito mortality (*µ*_*M*_) versus exposure coefficient demonstrating vector survival constraints. **(C)** Adult avian mortality (*µ*_*a*_) versus exposure coefficient showing demographic effects. **(D)** Mosquito biting rate (*a*) versus exposure coefficient revealing the primary transmission threshold. **(E)** Avian infectivity (*δ*) versus exposure coefficient displaying transmission efficiency boundaries. White lines indicate ℛ_0_ = 1 thresholds separating viable transmission from extinction.

Systematic parameter exploration confirms three distinct dynamical patterns predicted by ℛ_0_ analysis and stability theory. Fig. 9 illustrates these dynamics using biologically realistic parameter values, where outcomes are determined by ℛ_0_ and demographic stability characteristics.

**Fig. 9:**
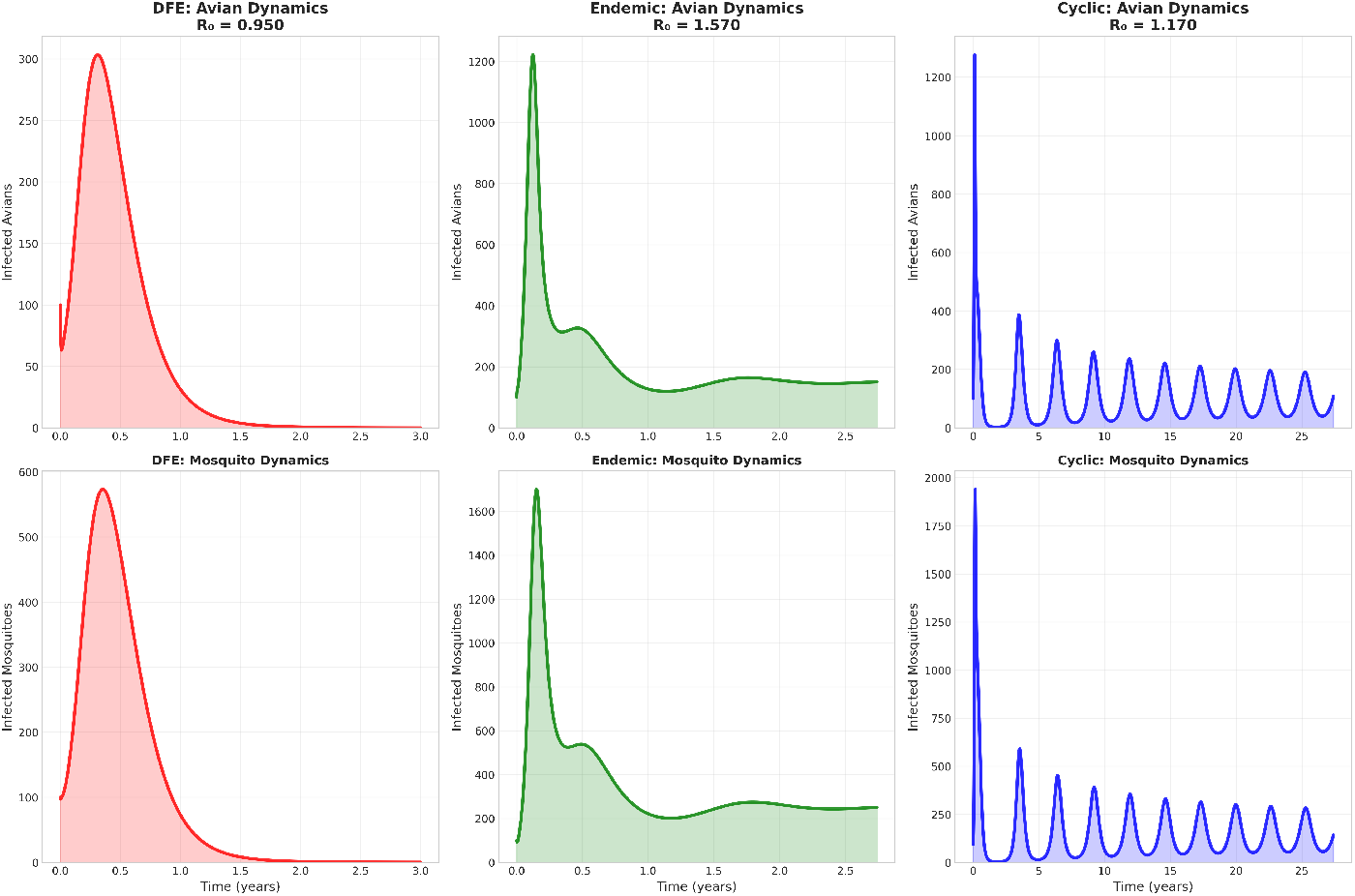
Population dynamics of total infected vectors/hosts across dynamical patterns. **Top row:** Total infected avians (filled curves) illustrating distinct dynamical patterns over extended temporal periods. **Bottom row:** Total infected mosquitoes (filled curves) showing corresponding vector dynamics. The three dynamical patterns illustrate fundamentally different long-term behaviors: extinction following initial outbreak (DFE), convergence to stable endemic levels (Stable EE), and persistent oscillations with 2–3 year periodicity (Cyclic/Unstable EE), consistent with observed EEEV outbreak patterns in nature.

### Disease-Free Equilibrium (ℛ_0_ = 0.950)

Parameter combinations with low mosquito biting rate (*a* = 0.8 day^−1^) and high mosquito mortality (*µ*_*M*_ = 0.18 day^−1^) produce realistic outbreak dynamics that peak around 5–6 months, and end up falling into complete extinction over 2–3 years. This pattern matches observed EEEV outbreaks in marginal habitats where environmental conditions prevent sustained transmission despite initial amplification [2, 26].

### Stable Endemic Equilibrium (ℛ_0_ = 1.570)

Balanced parameter values (*a* = 2.2 day^−1^, *α* = 0.5, *µ*_*M*_ = 0.08 day^−1^) result in stable transmission that rapidly converge to an equilibrium within 2–3 years. While this simulation demonstrates *standard* stable dynamical patterns, SHAP analyses revealed that *optimal* stable endemic conditions are characterized by higher mosquito biting rates and exposure coefficients combined with lower mosquito mortality. It is further verified that these balanced parameters better represent a standard rather than an optimal stable configuration, as seen in Fig. 6, since the stable EE (negative SHAP values) favors parameter combinations that result in enhanced transmission efficiency.

### Cyclic Dynamics (ℛ_0_ = 1.170)

Base parameter values from Table 1 (*ψ*_*B*_ = 0.05, *c* = 50,000, *a* = 2.5 day^−1^, *α* = 0.3) produce pronounced 2–3 year oscillatory behavior matching northeastern surveillance patterns. Despite having a moderate ℛ_0_, intermediate exposure coefficient values create demographic instability where preferential hatch-year exposure leads to periodic infection outbreaks followed by population recovery cycles. Consequently, this generates sustained oscillations rather than a stable endemic transmission.

### 5.1. Age-Structured Mechanisms

Detailed analysis of age-structured dynamics reveals the biological mechanisms underlying each dynamical pattern. Fig. 10 demonstrates how differential exposure between young and adult hosts drives system-level behavior, validating theoretical predictions about the critical role of host age-structure in EEEV transmission.

**Fig. 10:**
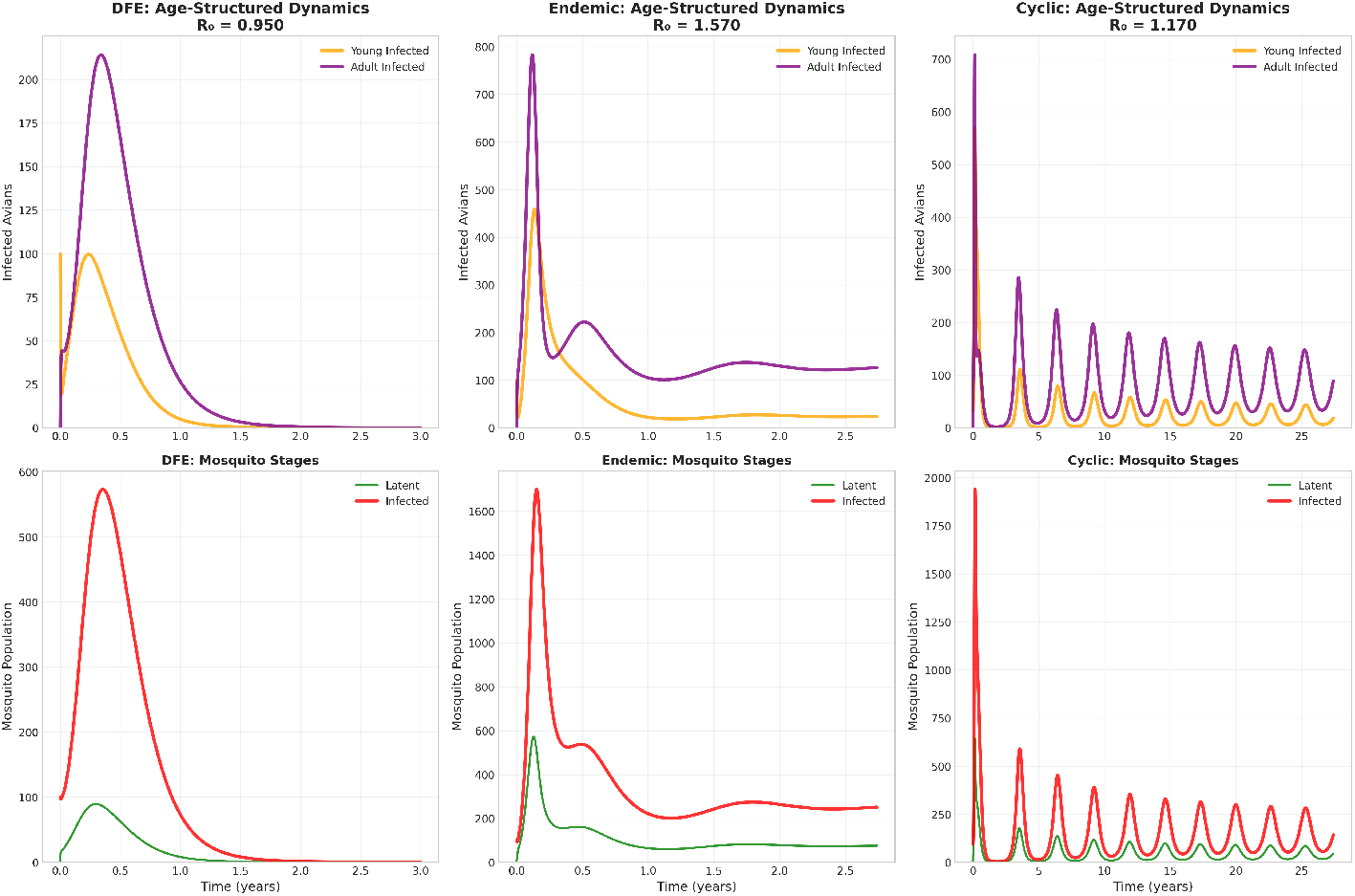
Age-structured population dynamics across dynamical patterns. **Top row:** Young (orange) and adult (purple) infected avian populations demonstrate differential age-structured responses. The disease-free equilibrium (ℛ_0_ = 0.950) shows rapid extinction after the initial outbreak around month 5–6. The stable EE (ℛ_0_ = 1.570) exhibits sustained transmission after the first peak with adults maintaining higher infection levels through time. Cyclic dynamics (ℛ_0_ = 1.170) display pronounced oscillations every 2–3 years, driven by age-structured amplification effects. **Bottom row:** Corresponding mosquito stage dynamics showing latent (green) and infected (red) populations, with cyclic dynamical patterns demonstrating synchronized periodic outbreaks in both compartments.

In the disease-free equilibrium, both age classes show a similar rapid convergence to extinction following initial exposure, confirming that ℛ_0_ < 1 prevents sustained transmission regardless of age-specific differences. Both age classes experience initial infection peaks, with adults reaching higher infection levels than young, before infections completely die out.

Stable endemic dynamics illustrate sustained transmission with adult avians facing higher infection levels than young avians due to longer lifespans and slower demographic turnover. This age-structured pattern emerges under balanced parameter conditions that avoid demographic oscillations seen in cyclic dynamics, consistent with SHAP analysis indicating higher *α* values favor transmission stability (Fig. 6). Cyclic dynamics reveal the mechanistic basis of oscillatory transmission through pronounced age-structured instability. Young avians reach peak infections of approximately 120 individuals during outbreak years before declining during inter-epidemic periods, while adult avians display larger amplitude oscillations (∼ 280 peak infections) with similar decline patterns. This demographic instability emerges when *α* = 0.3, indicating preferential exposure in young hosts, consistent with SHAP analysis identifying lower *α* values as drivers of cyclic behavior. The resulting oscillatory patterns exhibit biologically plausible periodic behavior with multi-year cycle characteristics. Mosquito infection dynamics exhibit fluctuations between epidemic peaks (∼ 600 peak infected vectors) and inter-epidemic minimums approaching near-extinction levels, reflecting the characteristic cyclical population dynamics of this transmission pattern.

These simulation results demonstrate strong correspondence with parameter importance rankings identified through sensitivity analysis and machine learning approaches. The mosquito biting rate (*a*) exhibits its sensitivity-predicted dominant influence by controlling transitions across the ℛ_0_ = 1 threshold that separates persistence from extinction. Meanwhile, the exposure coefficient (*α*) confirms its critical role in determining stability versus oscillatory dynamics above the same threshold. The quantitative agreement between theoretical predictions and simulated outcomes supports the EEEV age-structured model framework’s ability to capture essential biological mechanisms governing transmission. Distinct parameter combinations produce the three dynamical patterns at thresholds consistent with ℛ_0_ analysis and mechanistic pathways identified through SHAP analysis.

## 6. Discussion

Our integration of Sobol global sensitivity analysis, SHAP parameter importance, and dynamical simulations provides complementary insights into EEEV transmission mechanisms. Sobol analysis quantifies parameter influence on transmission potential, SHAP analysis identifies demographic drivers of endemic behavior, and model simulations validate theoretical predictions against observed patterns. This multi-method approach underscores the necessity of integrating diverse analytical perspectives to achieve a complete understanding of complex biological systems [31].

Sensitivity analyses present different parameter hierarchies that reveal distinct biological mechanisms. Sobol analysis ranks mosquito biting rate (*a*) as dominant (ST = 0.368–0.491), followed by avian infectivity (*δ*; ST = 0.120–0.231) and mosquito mortality (*µ*_*M*_ ; ST = 0.257). In contrast, SHAP analysis ranks adult avian mortality (*µ*_*a*_) as the most important parameter, followed by exposure coefficient (*α*), adult virulence (*ν*_*a*_), mosquito mortality (*µ*_*m*_), and mosquito biting rate (*a*). Remarkably, while *µ*_*a*_ achieves SHAP importance values up to twice those of the second most important parameter, it exhibits minimal Sobol sensitivity. This provides critical biological insight: while *µ*_*a*_ barely affects whether transmission can occur, it fundamentally determines whether endemic transmission exhibits stable or oscillatory dynamics within the viable parameter space.

The exposure coefficient (*α*) demonstrates consistent importance values across both analytical approaches and aligns well with simulation results. This convergence is evident across all analyses: Sobol analysis demonstrates *α*’s strong influence on ℛ_0_ (ST = 0.212), SHAP analysis (SHAP = 0.221) identifies threshold behavior at intermediate values (0.2–0.4) that generate demographic instability, and simulations confirm these predictions by reproducing cyclic dynamics exclusively within this critical range. The agreement between methods validates *α* as a key parameter, with simulations confirming that preferential exposure in young avians generates the demographic instability characteristic of northeastern EEEV patterns, while balanced exposure (*α* = 0.5) yields stable dynamics in both analytical frameworks.

Cyclic dynamics prove remarkably rare, occurring in only 1.5% of endemic equilibria across 100,000 parameter combinations analyzed using tree-based machine learning algorithms. Model simulations validate this finding by illustrating three distinct transmission patterns: disease-free equilibria/extinction (ℛ_0_ = 0.950), stable endemics (ℛ_0_ = 1.570), and cyclic dynamics (ℛ_0_ = 1.170). SHAP ridge plots (Fig. 7) reveal that cycles emerge through precise demographic balance rather than transmission optimization, with the threshold effects for both *µ*_*a*_ and *α* proving essential for generating oscillatory patterns reproduced in dynamical simulations.

The biological implications are clear from the methodological convergence seen between Sobol sensitivity, SHAP importance, and model simulations. Sobol analysis confirms that transmission viability depends on critical demographic thresholds, SHAP analysis reveals how specific age-structured exposure patterns generate instability, and simulations demonstrate that only rare parameter combinations reproduce the cyclic dynamics observed in northeastern EEEV surveillance data. Stable dynamics emerge when parameters optimize transmission and maintain demographic stability: high mosquito biting rates (*a*) and adult avian mortality (*µ*_*a*_), balanced exposure (*α* ≥ 0.5), and low mosquito mortality (*µ*_*M*_) and adult virulence (*ν*_*a*_). Unstable dynamics require the converse: demographic instability through low *a* and *µ*_*a*_, lower exposure to young avians (0.2 ≤ *α* ≤ 0.4), and high *µ*_*M*_ and *ν*_*a*_. Age-structured sensitivity patterns present further validation through simulation results, with young and adult avians showing differential responses to exposure and demographic parameters, reflected in the simulated oscillations where young avians peak at approximately 120 infections while adults peak at approximately 280 during cyclic outbreaks (Fig. 10).

This methodological integration advances EEEV disease modeling by demonstrating that traditional sensitivity analysis, machine learning approaches, and dynamical simulations address different but complementary biological questions. Our analytical approaches reveal critical threshold effects with direct intervention guidance. Within the critical exposure range (0.2 ≤ *α* ≤ 0.4), sensitivity analysis predicts—and simulations confirm—that minor changes in mosquito feeding behavior could dramatically shift regional transmission patterns from stable to oscillatory dynamics [7, 13].

## 7. Conclusions and Future Work

Our findings establish that EEEV transmission dynamics emerge from two distinct biological mechanisms: vector-host interaction parameters that determine outbreak potential and host demographic parameters that govern whether transmission exhibits stable or oscillatory dynamics. The identification of adult avian mortality as the dominant driver of cyclic dynamics, despite minimal influence on transmission thresholds, provides new insights into EEEV transmission mechanisms. Model simulations confirm this by showing how demographic instability within critical parameter ranges generates the 2–3 year cycles documented in northeastern surveillance data [2,13]. The methodological framework presented in this paper demonstrates that a comprehensive understanding of complex biological systems requires analytical approaches that capture both threshold effects and dynamical mechanisms.

This framework opens multiple avenues for advancing EEEV research across temporal, spatial, and environmental dimensions. Temporal extensions could incorporate seasonal variation in parameter importance analyses to reveal how demographic and transmission mechanisms shift throughout annual cycles, identifying optimal intervention timing. Spatial applications could extend SHAP analysis to geographically explicit models, explaining regional transmission differences and enabling the development of targeted management strategies. Environmental investigations could examine how climate change alters the identified parameter hierarchies, informing long-term surveillance planning, given the sensitivity of demographic stability parameters to environmental conditions. Beyond its application to EEEV, this multi-method approach could enhance understanding of other arboviral systems where demographic processes and transmission efficiency operate through distinct parameter sets [16, 29].

Our findings directly inform EEEV management by pinpointing adult host mortality and age-structured exposure as the fundamental drivers that determine cyclic versus stable transmission patterns. The dominance of these population-level parameters in determining transmission dynamics suggests that habitat conservation and close monitoring of host populations may be more effective than traditional vector control approaches. By demonstrating that EEEV cycles emerge from intrinsic host population dynamics rather than external drivers, this research redirects management paradigms toward population-based interventions and early warning systems grounded in the critical biological parameters identified here [10, 24].

## Acknowledgments

We would like to first acknowledge Drs. Kevin Caillouët and Christopher Kribs for reviewing the paper before publication for clarity, accuracy, and completeness. We would also like to thank Tarleton State University’s High Performance Computing Lab for allowing us to use their resources through the formative years of this research project. This work was partially funded by the Biotechnology Training Program (NIH NIGMS T32GM135066).

